# Alzheimer’s disease associations with increased Biondi body amyloid in hippocampal-associated choroid plexus epithelial cells and ependymal cells

**DOI:** 10.64898/2026.05.25.727667

**Authors:** Eunice Shin, Michelle Kim, Todd J. Soo, Olivia T. Espericueta, Erfan Zolfaghari, Michael J. Neel, Brett A. Johnson, Edwin S. Monuki

**Affiliations:** Department of Pathology & Laboratory Medicine, University of California Irvine, Irvine, CA, USA; Department of Developmental & Cell Biology, University of California Irvine, Irvine, CA, USA; Sue and Bill Gross Stem Cell Research Center, University of California Irvine, Irvine, CA, USA; Alzheimer’s Disease Research Center, University of California Irvine, Irvine, CA, USA

## Abstract

To resolve discrepancies in the literature regarding the association between Alzheimer’s disease (AD) and Biondi body (BB) amyloid in choroid plexus epithelial cells (CPECs), we investigated postmortem hippocampal paraffin blocks with and without a neuropathological diagnosis of AD (n=26-27 each). Similar to previous studies, age was associated with an increased fraction of hippocampal-associated CPECs bearing thioflavin S-positive BBs (p=0.004). In addition, we found that paraffin block storage time was associated with decreased BB detectability (p=0.038) while sex had no effect (p=0.577). Controlling for age, sex, and storage time, AD was associated with a near-significant increase in the BB-containing CPEC fraction (p=0.066) and a significantly greater load of BB-like amyloid in hippocampal-associated ependymal cells (p=0.032). The AD-BB association contrasts with our findings on choroid plexus from the atrium of the lateral ventricle, which lacked this association. We discuss potential explanations for the apparent discrepancy such as regional amyloid cross-seeding.

## INTRODUCTION

The choroid plexus (ChP) is a specialized tissue that plays a pivotal role in producing the cerebrospinal fluid (CSF). The ChP is located within each of the four ventricles of the brain, providing the CSF that protects and nourishes the central nervous system (CNS) while also removing waste and toxins (Lun et al., 2015; Katada et al., 2025). The ChP also establishes the blood-CSF barrier via choroid plexus epithelial cells (CPECs), which are interconnected by tight junctions (Laterra et al., 1999).

Biondi bodies (BBs) are intracellular lipofuscin-associated amyloid aggregates described by Biondi in the 1930s (Biondi, 1933). BB accumulation generally increases with age, becoming most prominent from middle age onward (Miklossy et al., 1998; Wen et al., 1999; Cottrell et al., 2001; Neel et al., manuscript in preparation). Recent findings from Ghetti et al. have further characterized their molecular composition, identifying lipofusin-like parts and the lysosomal membrane protein TMEM106B as a major constituent of BB amyloid (Ghetti et al., 2024).

Studies have reported elevated BB accumulation in the ChP of individuals with AD (Miklossy et al., 1998; Wen et al., 1999). However, the ventricular location of ChP examined was not specified in either study, and upon examining ChP from the atrium of the lateral ventricle, we found no association between BB accumulation and AD (Neel et al., manuscript in preparation). One possible explanation is that the other researchers examined ChP from another site, such as the temporal horn of the lateral ventricle. Temporal ChP is adjacent to the hippocampus, a structure central to memory formation, spatial navigation, and learning, which is also particularly vulnerable to the extracellular amyloid plaques and intracellular neurofibrillary tangles that are the hallmarks of AD neuropathology (Rao et al., 2022).

There has also been inconsistency and conflation in the literature concerning CPECs and ependymal cells on multiple levels, including with regard to BBs. Biondi originally described amyloid aggregates in both CPECs and ependymal cells, and thus both may be considered “Biondi bodies” (Biondi, 1933). In addition, the presence of TMEM106B in the amyloid inclusions of both CPECs and ependymal cells raises the possibility of a shared molecular composition (Ghetti et al., 2024). Miklossy et al. (1998) analyzed BBs in CPECs and ependymal cells collectively, noting an AD effect. It is important to note, however, that CPECs and ependymal cells have distinct developmental origins, trajectories, structures, and functions (Lehtinen et al., 2013, Yao et al., 2023; Groh et al., 2024), and the two cell types could certainly be differentially affected by AD.

The present paper addresses the apparent discrepancy and conflation in AD-BB associations by introducing region- and cell type-specific analyses of BB pathology. The first study examines BB prevalence in CPECs from the temporal horn of the lateral ventricle – hereafter referred to as “hippocampal-associated CPECs” – and tests whether these CPECs show a stronger AD effect than those from the atrium. The second study examines the AD effect in the amyloid aggregates of ependymal cells, which we refer to as “BB-like” to distinguish them from CPEC BBs. Potential associations of BB prevalence with age, sex, and tissue storage duration are also examined.

## MATERIALS AND METHODS

Paraffin-embedded hippocampal blocks that also included ChP were obtained from the UCI Alzheimer’s Disease Research Center (ADRC) tissue repository. The repository had over 100 AD cases and 29 non-AD control cases from autopsies between 1992 to 2008. Of the AD cases, 29 were selected for age and sex matching to the 29 control cases, yielding a total of 58 cases. Of these, 53 (Supplemental Table) had detectable ChP upon sectioning and were further analyzed. These samples were originally fixed, embedded in paraffin blocks, and sectioned at 5-µm thickness by the Experimental Tissue Resource core facility.

Slides were stained with primary antibodies targeting AE2 (anion exchanger protein; AE2 mouse monoclonal at 1:200; Santa Cruz Biotechnology SC376632) and AQP1 (aquaporin-1 water channel protein; AQP1 rabbit polyclonal at 1:1000; Millipore AB2219). As AQP1 lies on the apical membrane of CPECs and AE2 in the basolateral membrane, together these antibodies circumferentially label the CPEC plasma membrane. Slides were prepared by baking, followed by deparaffinization in xylenes and rehydration through descending diluted ethanol solutions. They were then stained with 0.5% Thioflavin-S (ThS; Millipore-Sigma T1892-25G) for 10 minutes, followed by two five-minute washes in 50% ethanol, and three five-minute washes in phosphate buffered saline, pH7.4 (PBS). Antigen retrieval was performed in a vegetable steamer using 10mM sodium citrate (pH 6). After using the blocking solution to prevent non-specific binding, primary antibodies were applied and incubated overnight at 4 degrees Celsius. Fluorescently-labeled secondary antibodies (Alexa 647 donkey anti-rabbit at 1:500 and anti-mouse at 1:200) were then applied. Whole slides were digitally scanned at 20X magnification using an Aperio Versa 2 scanner at the ADRC Neuropathology Core and converted into HDF5 files.

Due to an absence of nuclear staining by Hoechst in many of the older tissue sections, the AE2/AQP1 membrane labelling approach was used in place of the staining protocol previously used for the atrial ChP (Neel et al., manuscript in preparation). To assess whether this difference introduced systematic variability in BB and CPEC identification, we conducted a validation trial using 22 cases of atrial ChP, which were stained with both Hoechst and AE2/AQP1. Each case was then exported as two separate HDF5 files – one displaying ThS and Hoechest channels, and the other displaying ThS and AQP1/AE2 channels – with the non-displayed channel turned off (Supplemental Figure 1). Comparison of the two approaches revealed excellent correlation (linear regression r=0.96) and no systematic difference in the percentage of BB-containing CPECs between staining methods (Supplemental Figure 2).

### Study 1 (BB prevalence in hippocampal-associated CPECs)

All annotations by the three annotators (ES, MK, and BJ) were conducted blind to any demographic information, including AD status. BB-containing hippocampal-associated CPECs were manually counted using a custom software client (Neel et al., manuscript in preparation). The client divided whole slide images into smaller randomly-sampled tiles, then presented the tiles sequentially to annotators. As each tile was annotated, the client calculated a moving coefficient of error in real time. Like the atrial ChP study (Neel et al., manuscript in preparation), the pre-defined threshold for this moving coefficient of error was set at 0.05 for 10 consecutive tiles. Once this threshold was reached, the software signaled completion and generated the percentage of CPECs containing BBs. To reach the pre-defined threshold, between 292-7304 CPECs were annotated per case.

### Study 2 (BB-like load in hippocampal-associated ependyma)

The ependymal lining from each hippocampal-associated ChP case was captured at 20x magnification, using a Keyence BZ-X810 microscope, a monochromatic photodetector, a 20X objective (NA 0.75), and a GFP filter set.

Because individual ependymal cells were difficult to distinguish for cell-by-cell annotation, amyloid loads within the ependymal layer were quantified (i.e., the percentage of ependymal area that was positive for ThS). Each of three annotators imported images of the ependyma into ImageJ with Bio-Formats Importer plugin and converted these to gray scale. Individual channels were separated, and the relevant channel was convolved and despeckled twice to reduce background noise. A threshold was then applied and manually adjusted until the background signal of the tissue section was minimized while preserving the ependymal lining and ventricle. Final threshold value was then recorded, and the processed images were imported to QuPath with the threshold value applied. The ependymal lining was manually outlined as the region of interest (ROI). With the processed image, this annotation was applied, and thresholding was used aiming to isolate the ependymal amyloid structures (see Supplemental Figure 3). The percentage of affected ependyma area was calculated by dividing the total positive area by the total area of the ROI.

The median value across three annotators (ES, MK, and BJ) was used for statistical analysis to reduce inter-rater variability. ANCOVA analysis was performed to examine the relationship between percentage of affected area and AD status with covariates of age, sex, and storage duration.

## RESULTS

### Study 1 – BB prevalence in hippocampal-associated CPECs

When plotting the raw values with respect to age, the percentages of BB-containing hippocampal-associated CPECs were generally lower than those in atrial CPECs (see Figure 1). Our prior study of atrial ChP, which used more recently-acquired samples, did not show a consistent difference among locations within the lateral ventricle (Neel et al., manuscript in preparation). The hippocampal blocks for the current study had been stored for 15-30 years compared to 1-8 years for the atrial ChP blocks. Consequently, storage time was deemed to be an appropriate covariate, in addition to age and sex, in the ANCOVA analysis of BB-affected CPECs and AD status.

**Figure 1.**
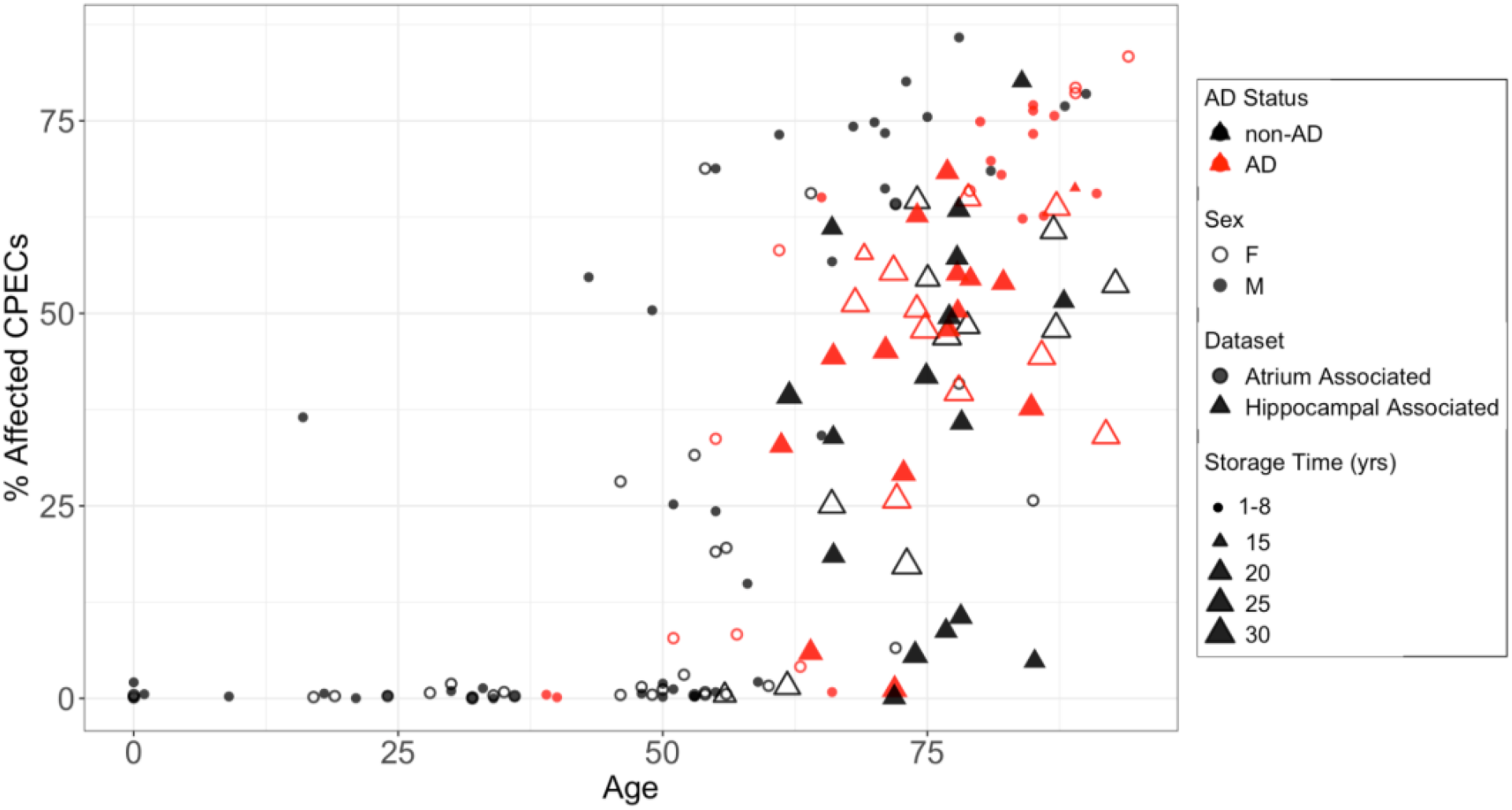
When viewed as raw data, the percentage of hippocampal-associated CPECs containing BBs (**triangles**) were consistently lower than the percentages in atrial CPECs from our prior study (**circles**). Hippocampal blocks were also stored for considerably longer than were the atrial ChP blocks (duration of storage is symbolized by the size of the symbols).

The ANCOVA analysis showed that AD status had a near-significant positive association with BB prevalence in CPECs, with AD cases showing a higher percentage of BB-affected CPECs compared to control cases (p=0.066, see Table 1). Age was a significant positive predictor of BB prevalence (p=0.004, see Table 1) consistent with previous studies. As we suspected, storage duration of the obtained tissue samples was a significant negative predictor (p=0.04, see Table 1), suggesting that longer storage was associated with reduced ThS signal. Sex was not a significant predictor (p=0.58, see Table 1). All effect plots are shown in Figure 2. The overall ANCOVA model was statistically significant, confirming that age and storage duration were necessary covariates.

**Figure 2.**
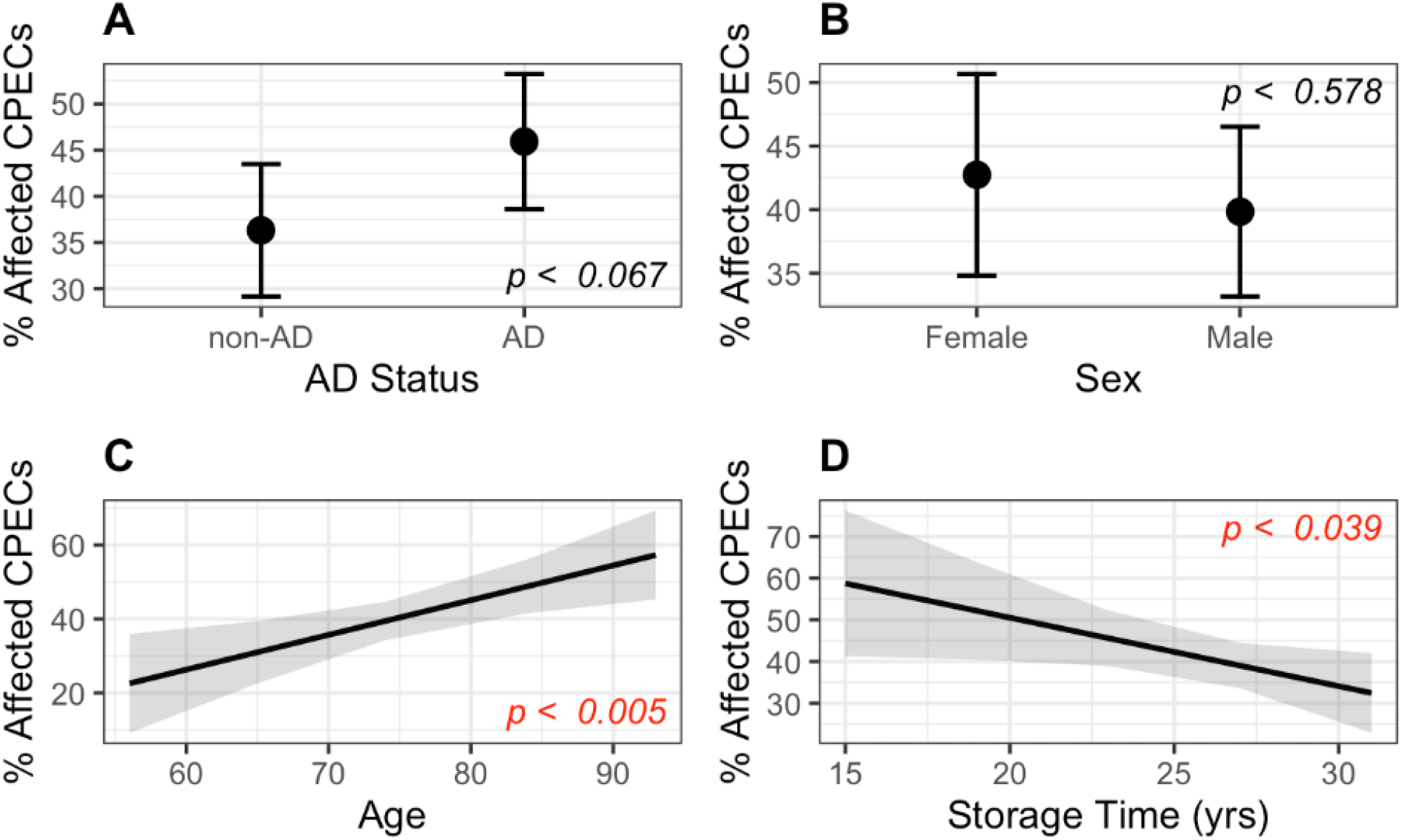
Effects of AD (**a**), sex (**b**), age (**c**), and storage time (**d**) on percentages of hippocampal-associated CPECs containing Biondi bodies, when corrected for other factors. AD was a near-significant factor, while age and storage time were significant at p<0.05.

**Table 1.**

| **ANCOVA Results: % Affected CPECs** |  |  |  |  |
| --- | --- | --- | --- | --- |
| F(4, 48) = 4.734, p = 0.003, R<sup>2</sup> = 0.283, Adjusted R<sup>2</sup> = 0.223 |  |  |  |  |
| **Variable** | **Estimate** | **Std. Error** | **t-value** | **p-value** |
| AD Status (AD vs. non-AD) | 9.594 | 5.104 | 1.880 | 0.066 <sup>†</sup> |
| Sex (Male vs. Female) | -2.899 | 5.164 | -0.561 | 0.577 |
| Storage Time | -1.644 | 0.773 | -2.128 | **0.038 \*** |
| Age | 0.939 | 0.313 | 2.998 | **0.004 \*\*** |
| <sup>†</sup> p < .10 * p < .05 ** p < .01 *** p < .001 |  |  |  |  |

### Study 2 – BB-like load in hippocampal-associated ependyma

As originally reported by Biondi (1933), BBs in CPECs had different morphologies compared to those in ependymal cells. CPEC BBs exhibited a wide range of highly-ordered morphologies such as rings and crescents (Neel et al., manuscript in preparation, and Supplemental Figure 3). In contrast, the amyloid inclusions in the ependyma appeared more frequently as wispy strands and small punctae.

As a measure of amyloid load, we determined the percent area of the ependyma stained by ThS. This percentage was significantly associated with AD status by ANCOVA with covariates of age, sex, and storage duration (p=0.03, see Table 2 and Figure 3). Age, sex, and storage duration were not significant predictors of ependymal amyloid load (p = 0.353, p = 0.300, and p = 0.235, respectively). Effect plots are shown in Figure 4. The overall ANCOVA model did not reach statistical significance, confirming that AD status emerged as a significant individual predictor, suggesting a specific association between AD and ependymal amyloid load independent of age, sex, and storage duration.

**Table 2.**

| **ANCOVA Results: % Affected Ependyma Area** |  |  |  |  |
| --- | --- | --- | --- | --- |
| F(4, 48) = 2.027, p = 0.106, R<sup>2</sup> = 0.147, Adjusted R<sup>2</sup> = 0.075 |  |  |  |  |
| **Variable** | **Estimate** | **Std. Error** | **t-value** | **p-value** |
| AD Status (AD vs. non-AD) | 3.044 | 1.378 | 2.210 | **0.032 \*** |
| Sex (Male vs. Female) | 1.457 | 1.390 | 1.048 | 0.3 |
| Storage Time | -0.249 | 0.207 | -1.202 | 0.235 |
| Age | 0.080 | 0.085 | 0.938 | 0.353 |
| <sup>†</sup> p < .10 * p < .05 ** p < .01 *** p < .001 |  |  |  |  |

**Figure 3.**
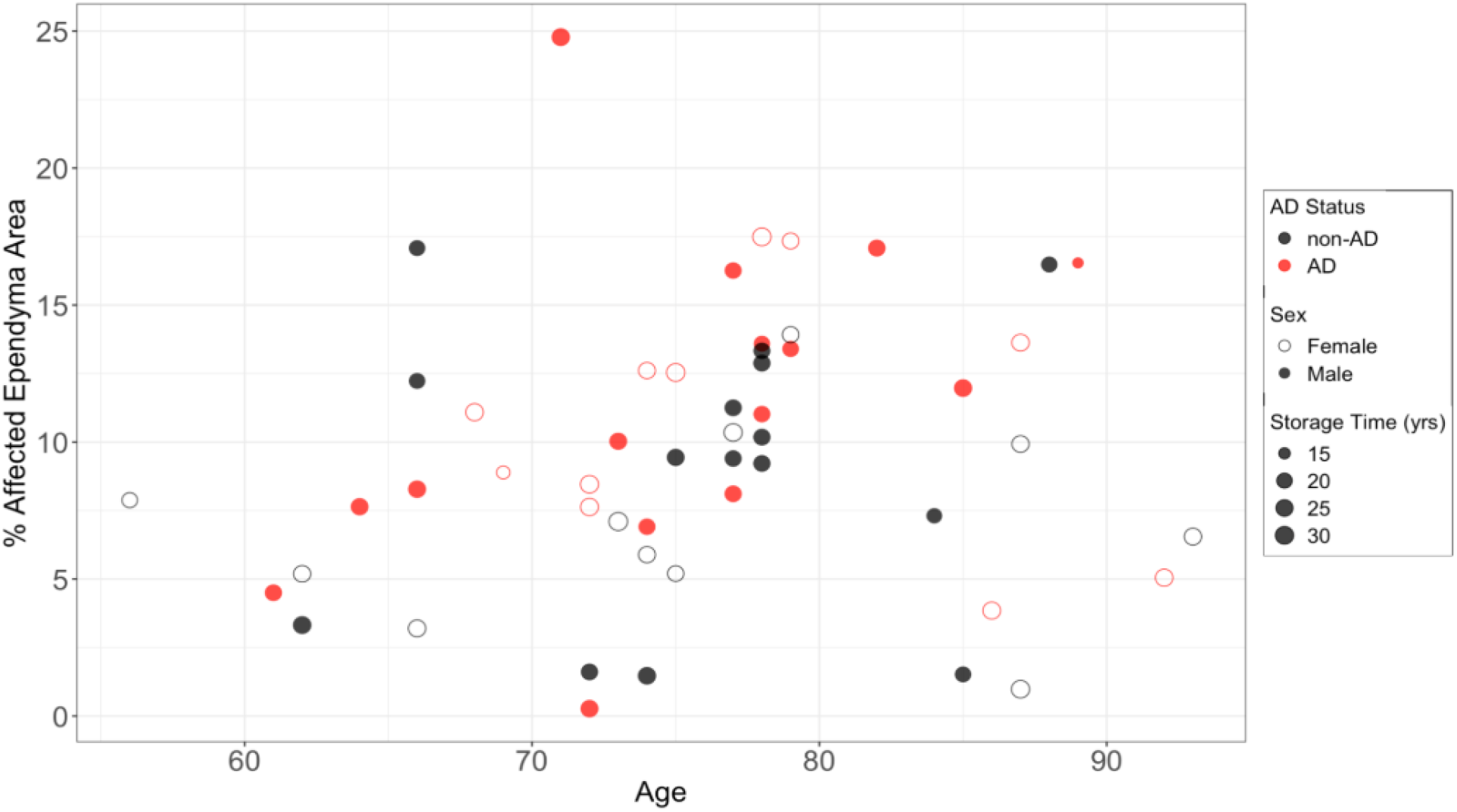
Raw data for the load of ependymal amyloid (percent area covered by ThS-staining) is expressed as a function of age.

**Figure 4.**
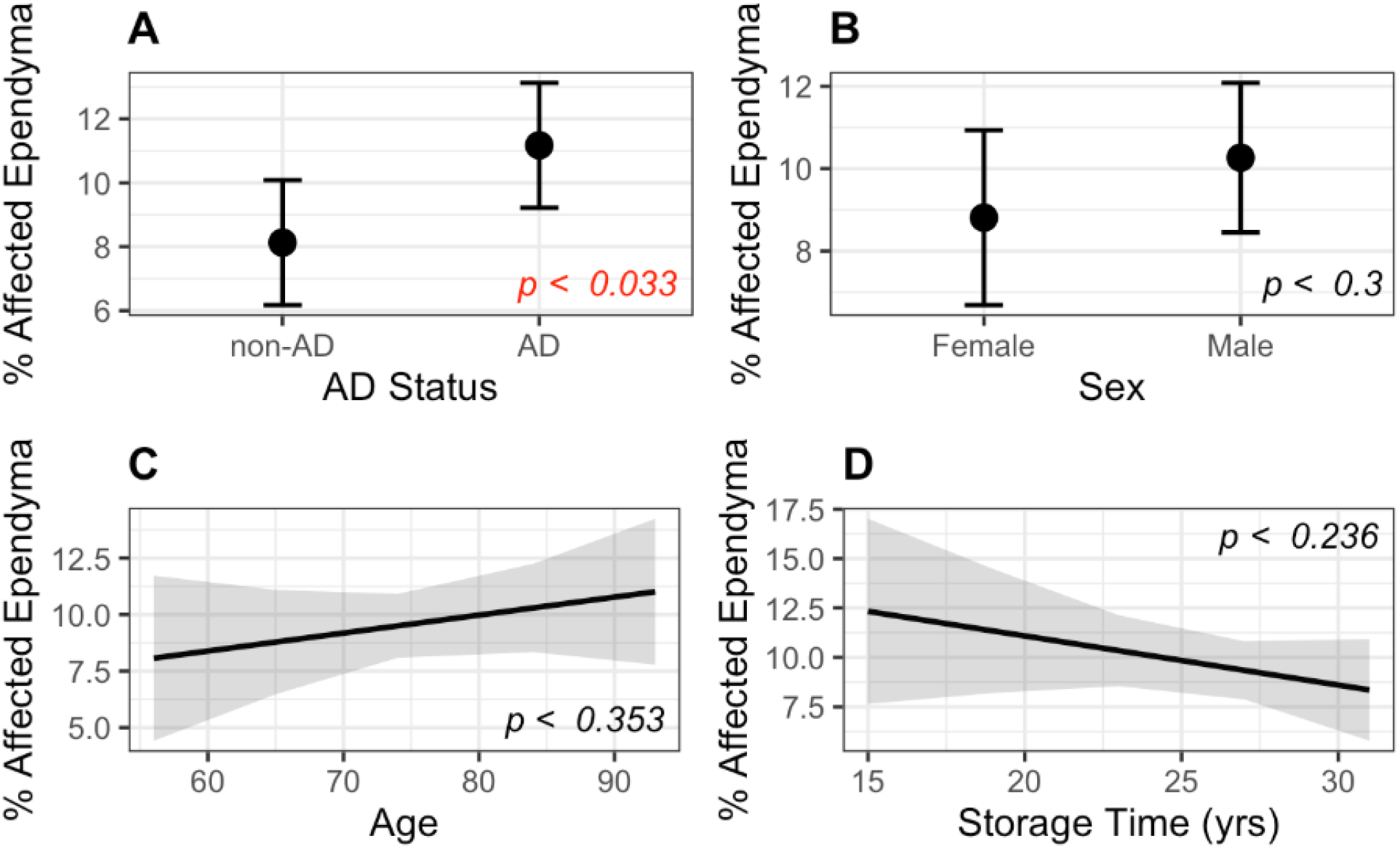
Effects of AD (**a**), sex (**b**), age (**c**), and storage time (**d**) on ependymal load when corrected for other factors. AD was a significant factor, while age and storage time displayed non-significant trends in the same directions (increases and decreases, respectively) that were seen for the hippocampal-associated CPEC Biondi bodies.

## DISCUSSION

In contrast to our prior findings of no AD effect on BB amyloid in atrial CPECs (Neel et al., manuscript in preparation), we now report a near-significant AD-associated increase in BBs in hippocampal-associated CPECs (p = 0.066). Additionally, we found an increased load of BB-like amyloid in the hippocampal-associated ependyma in the same tissue blocks (p = 0.032). These results may resolve the apparent discrepancy between our prior results and earlier reports of AD-associated increases in CPEC BBs, where details of ventricle locations were not provided (Miklossy et al., 1998; Wen et al., 1999) and where CPECs and ependymal cells were analyzed together (Miklossy et al., 1998).

The regional differences of AD effects on CPEC and ependymal amyloid suggests that proximity to the hippocampus may be an important factor in BB formation. The hippocampal parenchyma accumulates aggregates of hyperphosphorylated tau as well as plaques of amyloid-beta in association with AD (Rao et al., 2022), whereas parenchymal aggregates of TMEM106B – the amyloidogenic protein in BBs (Ghetti et al., 2024) – are associated with aging, but show no additional increase with AD (Schweighauser et al., 2022). The known cross-seeding among amyloid proteins (Ono et al., 2012; Vasconcelos et al., 2016) raises the possibility of tau and/or amyloid-beta cross-seeding of TMEM106B in neighboring ependymal cells and CPECs to produce BBs, or vice versa, given the relatively early age-specific prevalence of BBs in CPECs (Neel et al., manuscript in preparation).

The functional consequences of BB accumulation in CPECs and ependymal cells near the hippocampus remain to be established, but they may impair the many CPEC functions that support brain homeostasis (Lun et al., 2015; Katada et al., 2025). Likewise, amyloid accumulation may impair ependymal functions, including the movement of substances between CSF, interstitial brain fluid, and brain parenchyma. Thus, AD-associated BB formation may be part of a positive feedback process that further aggravates neurodegeneration.

## ACKNOWLEDGEMENTS

We thank the UCI neuropathologists and technicians at the Experimental Tissue Resource and UCI ADRC Neuropathology Core for their assistance in tissue procurement, processing, and slide scanning. We also thank Dan Hoang of the UCI ADRC for identifying AD cases that best matched the sex and age of available control hippocampal blocks. This work was supported by the following NIH awards: P50AG016573 and P30AG066519 (UCI ADRC), T32AG073088 and T32AG000096 (M.J.N.), and R21MH109036 and R21AG064640 (E.S.M.). This research would not have been possible without the generous donation by the tissue donors and their families.

---

**Supplemental Figure 1.**
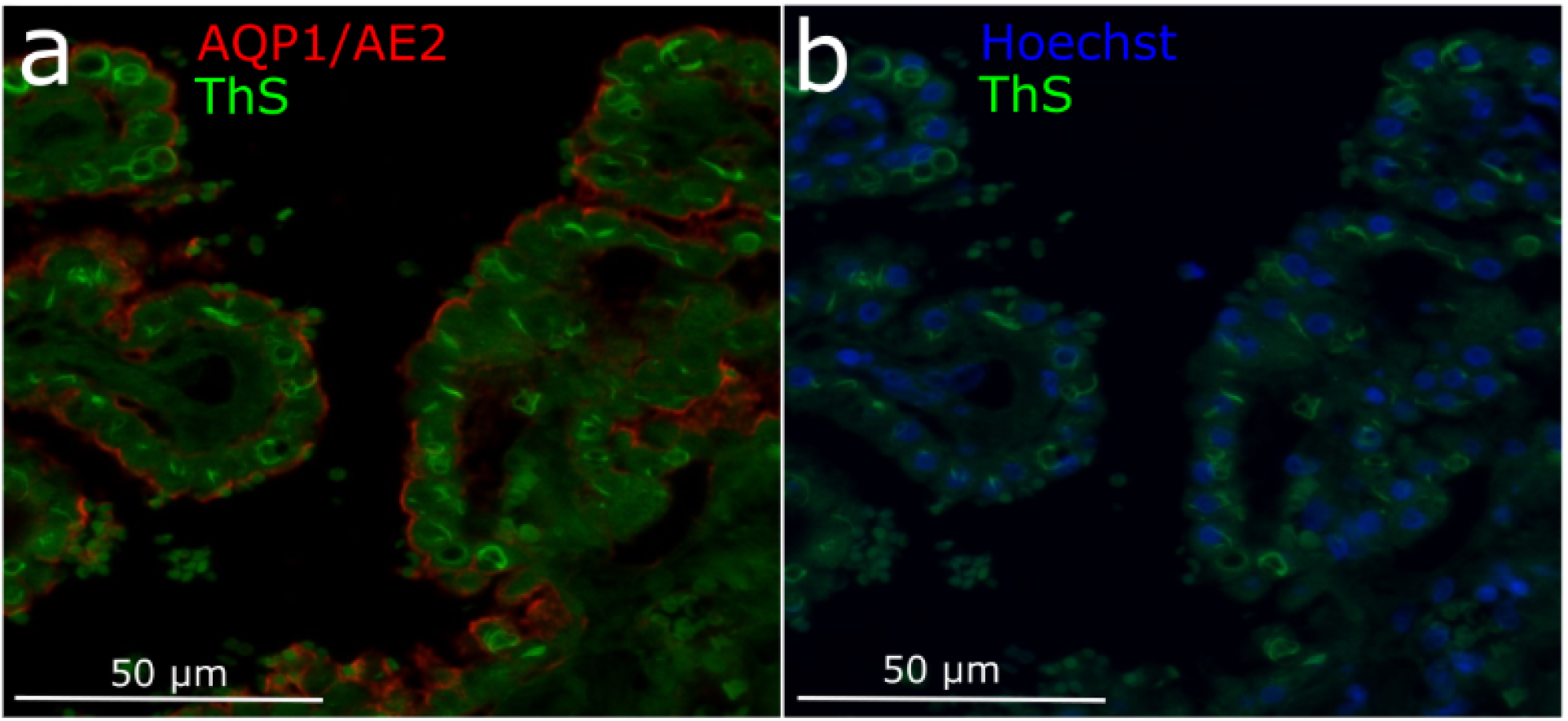
Images of the same region of a choroid plexus wherein individual CPECs were identified on the basis of membrane staining using immunofluorescence (**a**: AQP1, aquaporin 1; AE2, anion exchange protein 2) or Hoechst nuclear staining (**b**).

**Supplemental Figure 2.**
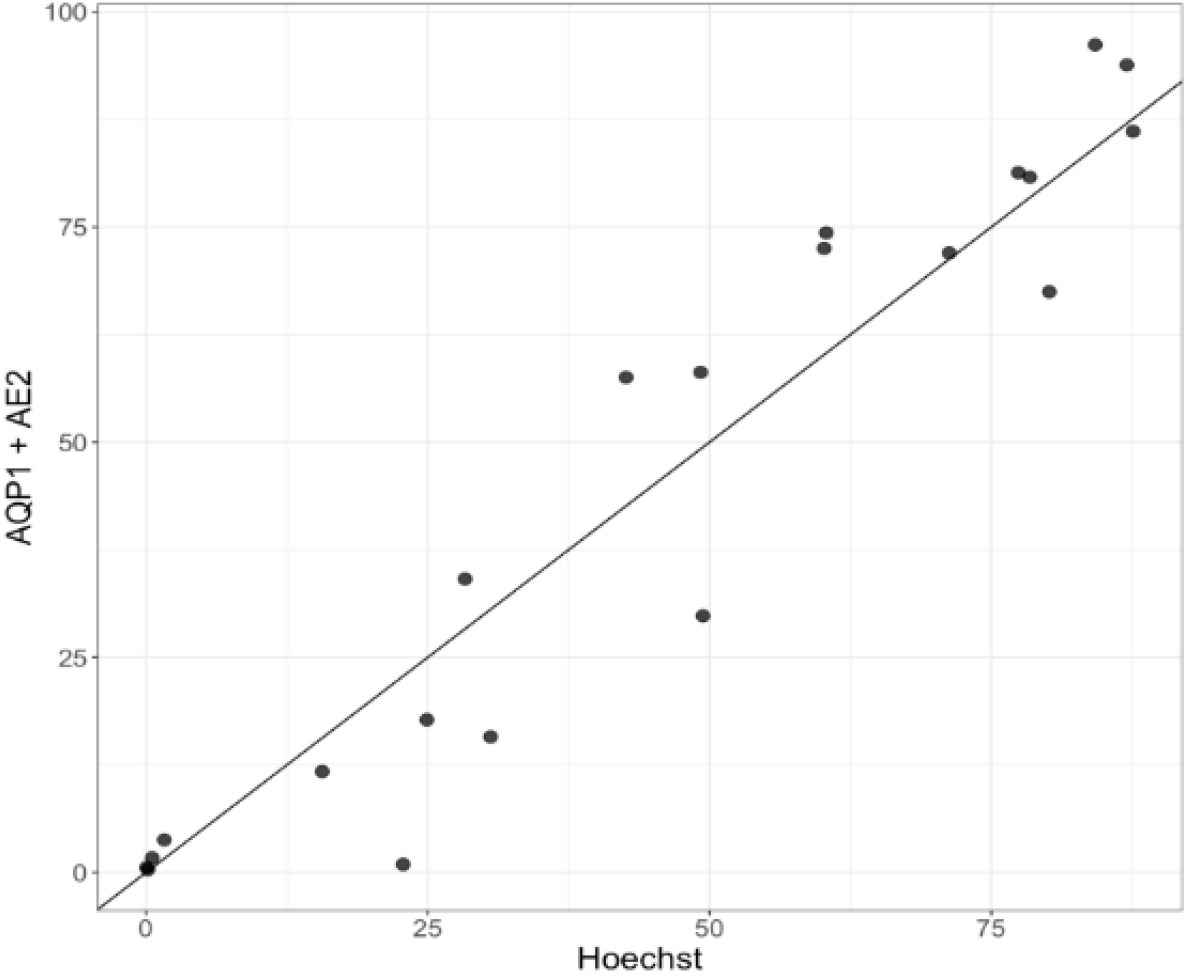
The two methods of identifying individual CPECs result in similar estimates of the percentage of CPECs containing BBs. The trendline is a result of linear regression, with an r value of 0.96.

**Supplemental Figure 3.**
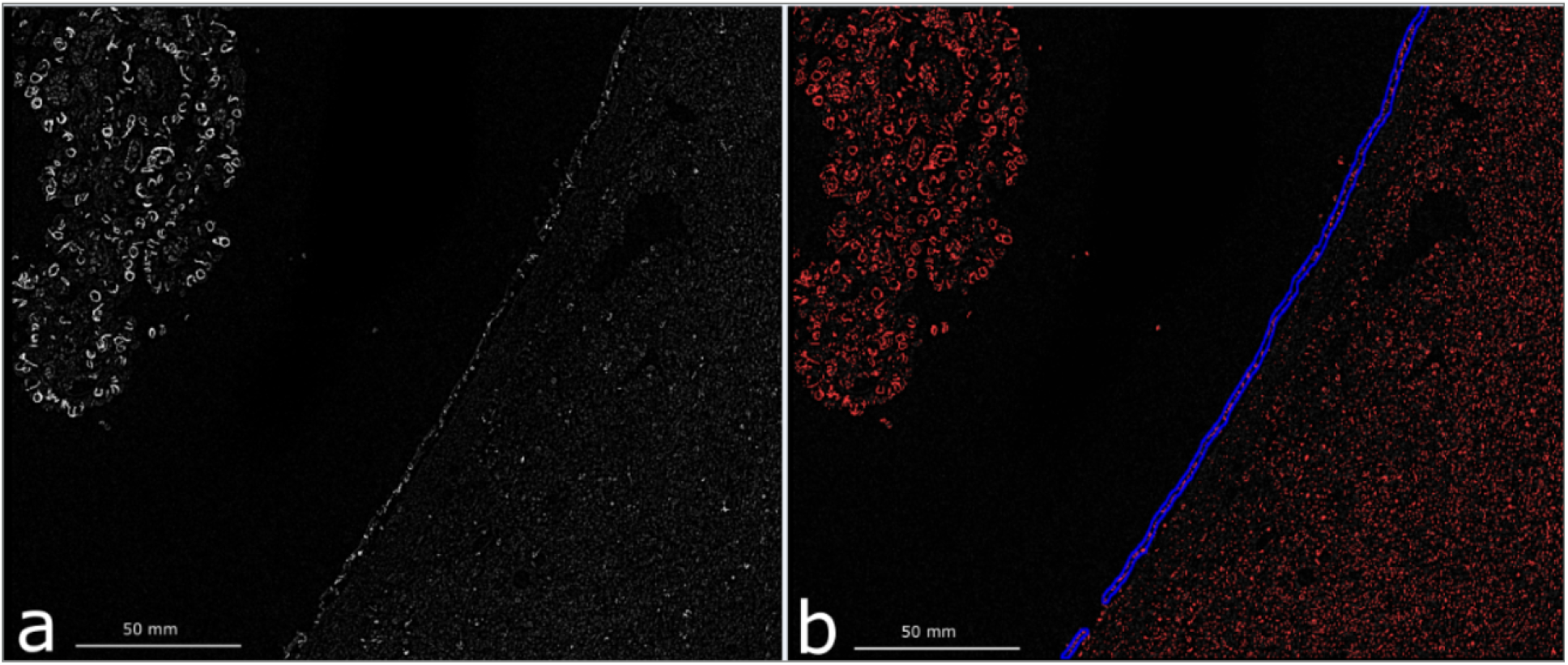
To determine the load of ependymal amyloid as a percentage of ependymal area stained by ThS, images were threshholded in ImageJ (**a**), the ependymal layers were outlined in QuPath (**b**), and the area covered by thresholded pixels within the outline were expressed as a percentage of that area.

